# Genetic basis and timing of a major mating system shift in *Capsella*

**DOI:** 10.1101/425389

**Authors:** Jörg A. Bachmann, Andrew Tedder, Benjamin Laenen, Marco Fracassetti, Aurélie Désamoré, Clément Lafon-Placette, Kim A. Steige, Caroline Callot, William Marande, Barbara Neuffer, Hélène Bergès, Claudia Köhler, Vincent Castric, Tanja Slotte

## Abstract

Shifts from outcrossing to self-fertilisation have occurred repeatedly in many different lineages of flowering plants, and often involve the breakdown of genetic outcrossing mechanisms. In the Brassicaceae, self-incompatibility (SI) allows plants to ensure outcrossing by recognition and rejection of self-pollen on the stigma. This occurs through the interaction of female and male specificity components, consisting of a pistil based receptor and a pollen-coat protein, both of which are encoded by tightly linked genes at the *S*-locus. When benefits of selfing are higher than costs of inbreeding, theory predicts that loss-of-function mutations in the male (pollen) SI component should be favoured, especially if they are dominant. However, it remains unclear whether mutations in the male component of SI are predominantly responsible for shifts to self-compatibility, and testing this prediction has been difficult due to the challenges of sequencing the highly polymorphic and repetitive ~100 kbp *S*-locus. The crucifer genus *Capsella* offers an excellent opportunity to study multiple transitions from outcrossing to self-fertilization, but so far, little is known about the genetic basis and timing of loss of SI in the self-fertilizing diploid *Capsella orientalis*. Here, we show that loss of SI in *C. orientalis* occurred within the past 2.6 Mya and maps as a dominant trait to the *S*-locus. Using targeted long-read sequencing of multiple complete S-haplotypes, we identify a frameshift deletion in the male specificity gene *SCR* that is fixed in *C. orientalis*, and we confirm loss of male SI specificity. We further analyze RNA sequencing data to identify a conserved, *S*-linked small RNA (sRNA) that is predicted to cause dominance of self-compatibility. Our results suggest that degeneration of pollen SI specificity in dominant *S*-alleles is important for shifts to self-fertilization in the Brassicaceae.

**Author Summary:** Already Darwin was fascinated by the widely varying modes of plant reproduction. The shift from outcrossing to self-fertilization is considered one of the most frequent evolutionary transitions in flowering plants, yet we still know little about the genetic basis of these shifts. In the Brassicaceae, outcrossing is enforced by a self-incompatibility (SI) system that enables the recognition and rejection of self pollen. This occurs through the action of two tightly linked genes at the *S*-locus, that encode a receptor protein located on the stigma (female component) and a pollen ligand protein (male component), respectively. Nevertheless, SI has frequently been lost, and theory predicts that mutations in the male component should have an advantage during the loss of SI, especially if they are dominant. To test this hypothesis, we mapped the loss of SI in a selfing species from the genus *Capsella*, a model system for evolutionary genomics. We found that loss of SI mapped to the *S*-locus, which harbored a dominant loss-of-function mutation in the male SI protein, and as expected, we found that male specificity was indeed lost in *C. orientalis*. Our results suggest that transitions to selfing often involve parallel genetic changes.

## Introduction

The shift from outcrossing to self-fertilization is one of the most common evolutionary transitions in flowering plants (Darwin 1876; Wright et al. 2013). This transition is favored when the benefits of reproductive assurance (Darwin 1876; Pannell and Barrett 1998; Eckert et al. 2006) and the transmission advantage of selfing (Fisher 1941) outweigh the cost of inbreeding depression (Charlesworth 2006).

The transition to self-fertilization often involves breakdown of self-incompatibility (SI). SI systems allow plants to recognize and reject self pollen through the action of male and female specificity components and modifier loci (Takayama and Isogai 2005). In the Brassicaceae, where the molecular basis of SI is particularly well characterized, SI is controlled by two tightly linked genes at the *S*-locus, *SRK* and *SCR*, which encode the female and male SI specificity determinants, respectively (de Nettancourt 2001). SRK is a transmembrane serine-threonine receptor kinase located on the stigma surface (Stein et al. 1991; Stein et al. 1996), and SCR is a small cysteine-rich protein deposited on the pollen coat, that acts as a ligand to the SRK receptor (Schopfer et al. 1999; Takayama et al. 2001). Direct interaction between SRK and SCR from the same *S*-haplotype results in inhibition of pollen germination (Takasaki et al. 2000; Takayama et al. 2001; Ma et al. 2016) through a signaling cascade involving several proteins (Nasrallah and Nasrallah 2014). This prevents close inbreeding and promotes outcrossing. At the *S*-locus, recombination is suppressed and rare allele advantage maintains alleles with different specificities (Wright 1939; Castric and Vekemans 2004; Vekemans et al. 2014), such that SI populations often harbor dozens of highly diverged *S*-haplotypes (Mable et al. 2003; Guo et al. 2009). In the sporophytic Brassicaceae SI system, expression of a single *S*-specificity provides greater compatibility with other individuals (Schoen and Busch 2009) and therefore *S*-haplotypes often form a dominance hierarchy, that determines which specificity is expressed in *S*-heterozygotes (Durand et al. 2014). At the pollen level, dominance is governed by dominance modifiers in the form of sRNAs expressed by dominant alleles that target sequence motifs specific to recessive alleles of *SCR*, resulting in their transcriptional silencing (Tarutani et al. 2010; Durand et al. 2014).

Despite the advantages of outcrossing, SI has been lost repeatedly in many different lineages, and there is a strong theoretical and empirical interest in the role of parallel molecular changes for repeated shifts to self-compatibility (SC) (Vekemans et al. 2014; Shimizu and Tsuchimatsu 2015). While the numerous genes that act as unlinked modifiers of SI potentially constitute a larger mutational target, theory predicts that mutations that result in degeneration of components of the *S*-locus itself should have an advantage (Porcher and Lande 2005). Theory further predicts that the probability of spread of mutations disrupting SI depends on whether they affect male or female functions, or both functions jointly (Charlesworth and Charlesworth 1979). In particular, mutations that disrupt male specificity should have an advantage over those mutations that disrupt female specificity, because male specificity mutations can spread faster through both pollen and seeds (Uyenoyama et al. 2001; Tsuchimatsu and Shimizu 2013). Finally, while dominant advantageous mutations should have a higher fixation probability in outcrossers, as expected from Haldane’s sieve (Haldane 1927), dominant *S*-alleles typically have low population frequencies (Llaurens et al. 2008), resulting in a lower probability that SC mutations occur on dominant than on recessive alleles. While degeneration of male specificity has contributed to loss of SI in several Brassicaceae species (Tsuchimatsu et al. 2010; Tsuchimatsu et al. 2012; Chantha et al. 2013; Shimizu and Tsuchimatsu 2015), it is unclear how general this pattern is, and few empirical studies have examined the contribution of dominant *S*-haplotypes to the loss of SI.

To test these hypotheses identification of causal mutations is required, a task that is challenging due to the high level of divergence among *S*-haplotypes with different specificities. One solution is to contrast functional and non-functional *S*-haplotypes that belong to the same *S*-haplogroup and ancestrally shared the same SI specificity. It has previously been difficult to obtain full-length sequences of the up to 110 kb long, highly polymorphic and repetitive *S*-locus, however thanks to the advent of long-read sequencing contiguous *S*-haplotypes can now be assembled with low error rates (Bachmann et al. 2018).

The crucifer genus *Capsella* is an emerging model for genomic studies of plant mating system evolution. In *Capsella*, SI is the ancestral state, as there is trans-specific shared *S*-locus polymorphism between the outcrossing SI species *Capsella grandiflora* and outcrossing SI *Arabidopsis* species (Guo et al. 2009). Nevertheless, SC has evolved repeatedly, resulting in two self-compatible and highly selfing diploid species, *Capsella rubella* and *Capsella orientalis*, as well as the selfing allotetraploid *Capsella bursa-pastoris*, which formed by hybridization and genome duplication between *C. orientalis* and *C. grandiflora* (Douglas et al. 2015). These species also differ greatly in their geographical distributions, with *C. bursa-pastoris* having a nearly worldwide distribution, whereas *C. rubella* is mainly found in Central and Southern Europe, and *C. orientalis* has a distribution ranging from Eastern Europe to Central Asia (Hurka et al. 2012). Finally, the SI outcrosser *C. grandiflora* is limited to northwestern Greece and Albania (Hurka et al. 2012).

When studying the consequences of selfing it is essential to distinguish between changes that occurred before and after the mating system shift. Understanding when and how SI was lost is thus crucial. In *C. rubella*, the transition to selfing has been intensely studied (Foxe et al. 2009; Guo et al. 2009; Brandvain et al. 2013; Slotte et al. 2013) and involved the fixation of a relatively dominant *S*-haplotype (Guo et al. 2009; Paetsch et al. 2010) most likely within the past 50-100 kya (Foxe et al. 2009; Slotte et al. 2013). Knowledge on the mode, timing and demographics of the transition to selfing in *C. rubella* has provided an evolutionary context for the study of genomic (Brandvain et al. 2013; Slotte et al. 2013), regulatory (Steige et al. 2015) and phenotypic (Slotte et al. 2012; Sicard et al. 2016) consequences of selfing. In contrast, we know little about the genetic basis and timing of loss of SI and transition to selfing in *C. orientalis*, although such information is important for proper interpretation of genomic studies of the effects of selfing and can provide general insights into the role of parallel molecular changes for convergent loss of SI.

Here, we therefore combined genetic mapping, long-read sequencing of *S*-haplotypes, controlled crosses, population genomic and expression analyses to investigate the loss of SI in *C. orientalis*, with the specific aims to: 1) test whether loss of SI maps to the *S*-locus, 2) identify candidate causal mutations for the loss of SI, 3) investigate the role of sRNA-based dominance modifiers, and 4) estimate the timing of loss of SI in *C. orientalis*. Our results are important for an improved understanding of the role of parallel molecular changes for transitions to selfing.

## Results

### SC maps to the S-locus as a dominant trait

We first asked whether loss of SI in *C. orientalis* maps to the canonical Brassicaceae *S*-locus. We therefore generated an F2 mapping population by crossing *C. orientalis* to a SI *C. grandiflora* accession. Interspecific F1 individuals were SC, indicating that SC is dominant. Our F2 mapping population segregated for SC, and we detected a single, significant (P<0.001) quantitative trait locus (QTL) for this trait, based on 304 F2 individuals genotyped at 549 markers (fig. 1A). The credible interval for this QTL includes the *S*-locus on chromosome 7 (fig. 1A), and SC was dominant over SI (fig. 1B). SC in *C. orientalis* thus maps as a dominant trait to a region encompassing the *S*-locus.

**Figure 1.**
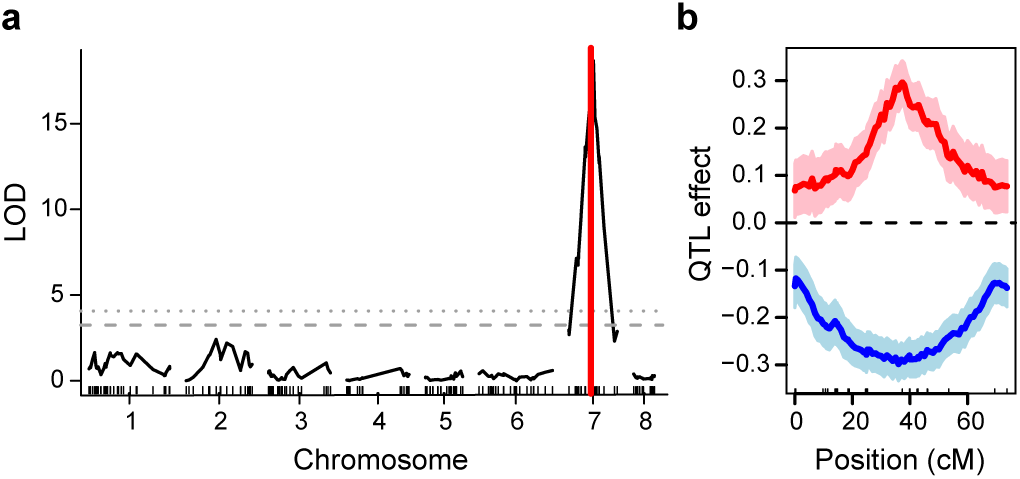
Self-compatibility is dominant and maps to the *S*-locus. **a.**Logarithm of odds (LOD) profile resulting from interval mapping of self-compatibility in an interspecific *C. orientalis* × *C. grandiflora* F2 population. The dotted and dashed lines indicates the 1% vs. 5% genome-wide permutation-based significance threshold. The red vertical line shows the location of the canonical Brassicaceae *S*-locus. The 1.5-LOD confidence interval ranges from position 6,241,223 to 8,742,368, whereas the *S*-locus is located between positions 7,523,602 and 7,562,919 on chromosome 7. **b.** Estimated quantitative trait locus (QTL) additive effect (red line) and dominance deviation (blue line) across chromosome 7. Light shaded regions indicate standard errors.

### Sequencing the S-haplotype of C. orientalis and a highly similar but functional S-haplotype from C. grandiflora

We next sought to identify candidate causal loss-of-function mutations at the *C. orientalis S*-locus. For this purpose, we first assembled full-length *S*-haplotype sequences of two *C. orientalis* accessions based on long-read sequencing of BACs (supplementary tables S1 and S2, Supplementary Material). To facilitate identification of candidate mutations for the loss of SI, we identified and sequenced a functional *C. grandiflora S*-haplotype (for details, see Materials and Methods), which had 98.3% protein sequence identity at *SRK* to that of *C. orientalis* (fig. 2A-C, supplementary table S3, Supplementary Material) and is likely to represent the same SI specificity based on criteria used in outcrossing *Arabidopsis* species (Castric et al. 2008; Tsuchimatsu et al. 2012). This *C. grandiflora S*-haplotype is also similar (93.4% protein sequence identity at *SRK*) to the functional *Arabidopsis halleri S12* haplotype (Durand et al. 2014) (fig. 2A-B, supplementary fig. S1, Supplementary Material), and hereafter we therefore designate it *CgS12*. We verified that *C. grandiflora* individuals with *CgS12* expressed *CgSCR12* and were SI by scoring pollen tube germination after controlled self-pollination (fig. 3, supplementary table S4, fig. S2-S4, Supplementary Material).

**Figure 2.**
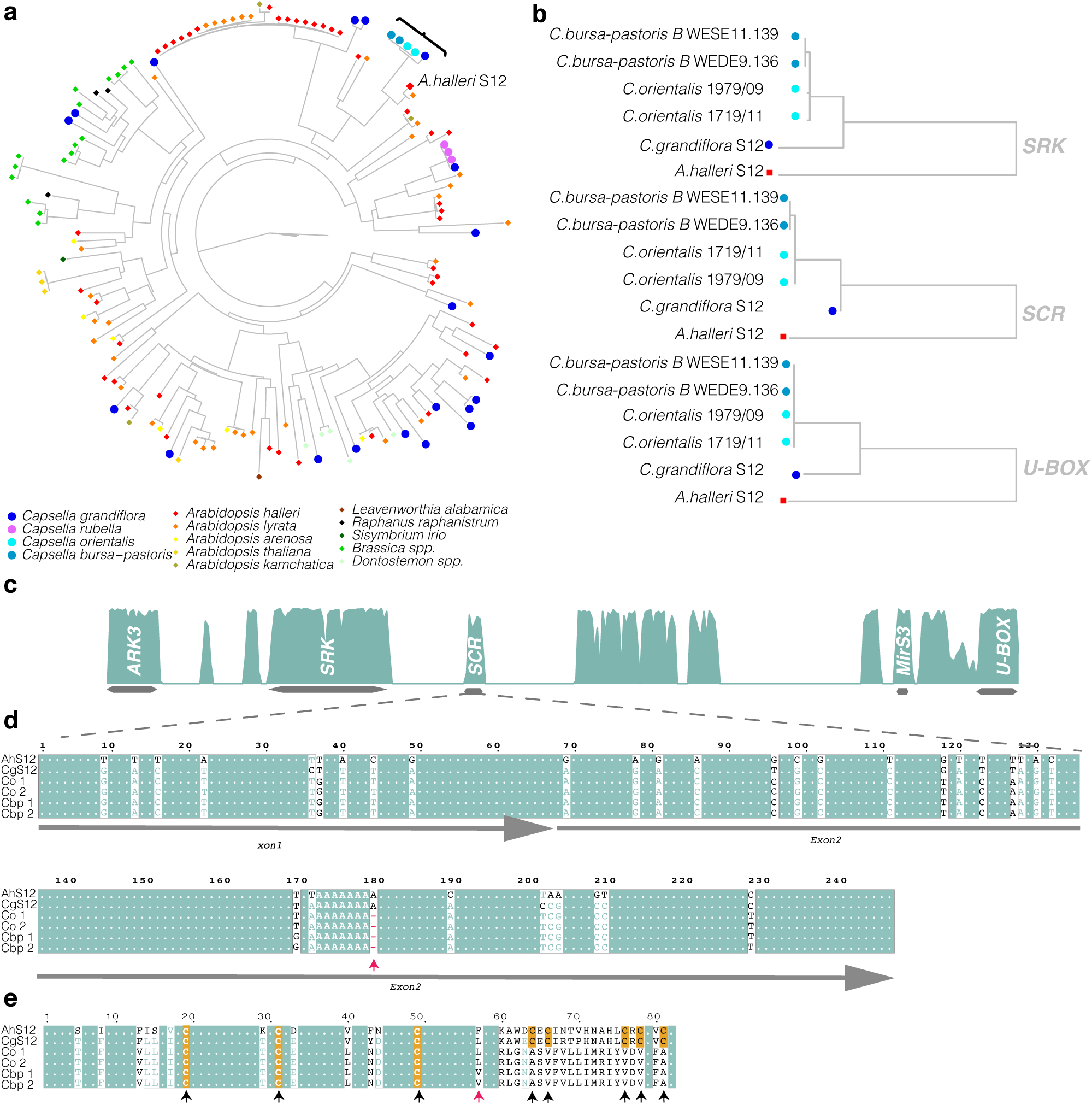
Sequence comparison of full-length *S*-haplotype sequences results in identification of a frameshift deletion in *C. orientalis SCR*. **a.**Phylogram of *SRK* sequences, showing the diversity of *S*-alleles among Brassicaceae and the close similarity of *SRK* from *A. halleri S12, C. grandiflora CgS12, C. orientalis* and *C. bursa-pastoris* (B subgenome). **b.**Maximum likelihood gene trees for three *S*-locus genes: *SRK, SCR* and *U-BOX* showing the relationship between *A. halleri S12, C. grandiflora CgS12, C. orientalis* and *C. bursa-pastoris* (B subgenome). **c.**Percentage of sequence similarity between *C. grandiflora CgS12 and C.orientalis* 1979/09 *S*-haplotypes. Gene position are indicated by grey bars. **d.**Alignment of *SCR* sequences from *A. halleri S12, C. grandiflora CgS12, C. orientalis* and *C.bursa-pastoris* (*B* subgenome) S-haplotypes. A frameshift deletion in the coding sequence (marked by a red arrow) is found in *C. orientalis* but not in the two SI species *A. halleri* and *C. grandiflora*. **c.** Predicted SCR amino acid sequences for *A. halleri S12, C. grandiflora CgS12, C. orientalis* and *C. bursa-pastoris* (*B* subgenome). The predicted protein sequence of *C. orientalis* lacks five conserved cysteine residues (indicated by black arrows and orange boxes). The position of the frameshift deletion is marked by a red arrow.

### A frameshift deletion in the male specificity gene SCR is fixed in C. orientalis

By comparing *S*-haplotype sequences from *C. orientalis* (SC) to *C. grandiflora CgS12* and *A. halleri S12* (both SI), we identified a single-base frameshift deletion in the *SCR* coding sequence of *C. orientalis* (fig. 2D). This frameshift is predicted to result in loss of 5 out of 8 conserved cysteine residues essential to the function of SCR (fig. 2E), likely resulting in loss of male specificity. To assess whether the deletion was fixed in *C. orientalis*, as we would expect for mutations that spread early during the transition to selfing, we analyzed whole-genome resequencing data (table S1, Supplementary Material) from additional *C. orientalis* accessions (table S1, Supplementary Material). We found that the *SCR* frameshift deletion was fixed across 32 samples of *C. orientalis* from 18 populations, consistent with expectations if the deletion was fixed in association with the loss of SI. The same deletion was found in *SCR* of the *C. bursa-pastoris* B subgenome, which is derived from *C. orientalis* (fig. 2D, fig. 2E). This finding is consistent with our previous inference that *C. orientalis* was selfing when it contributed to the origin of the allotetraploid *C. bursa-pastoris* (Douglas et al. 2015).

### Assessment of SI specificity

To assess whether male SI specificity is degenerated in *C. orientalis*, as we expect if SCR is nonfunctional, we crossed *C. orientalis* to *C. grandiflora* individuals harboring *CgS12*, which likely ancestrally shared the same SI specificity (fig. 2). As expected if the frameshift deletion impaired the function of *SCR*, pollen from *C. orientalis* successfully germinated on the stigma of *C. grandiflora* individuals harboring *CgS12* (fig. 3, fig. S2-S3, Supplementary Material). However, we also found evidence for degeneration of female specificity in *C. orientalis*, as pollen from *C. grandiflora* harboring *CgS12* germinated on the *C. orientalis* stigma (fig. S3-S4, Supplementary Material). In contrast to *SCR* however, we observed no major loss-of-function mutations in *C. orientalis SRK* or at the *S*-linked *U-box* gene, which may modify the female SI response (Liu et al. 2007). *SRK, U-box* and *SCR* are all expressed in flower buds of *C. orientalis* (table S4; fig. S4, Supplementary Material) and we currently cannot rule out that more subtle changes to their sequence or expression affect their function.

**Figure 3.**
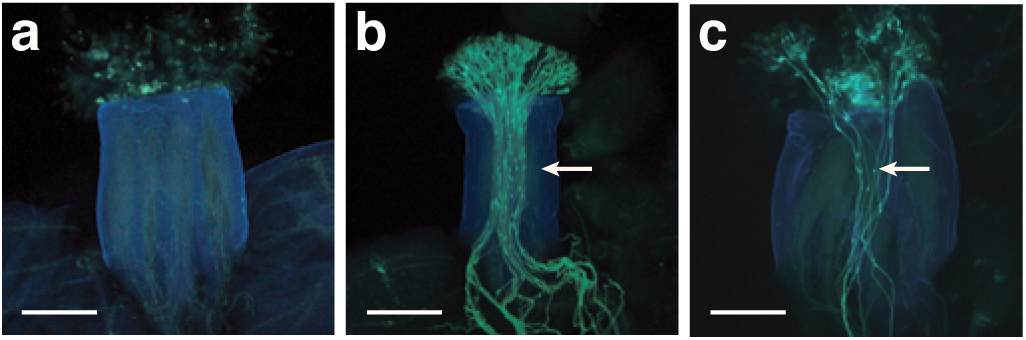
Male self-incompatibility specificity is disrupted in *C. orientalis*. **a.**Self-pollination of *C. grandiflora* carrying *CgS12* allele results in no pollen tube growth (incompatible reaction), demonstrating functional self-incompatibility. **b.**Pollination of *C. grandiflora* carrying *CgS12* with pollen from an individual carrying different *S*-haplotypes results in pollen tube growth (compatible reaction). **c.**Pollination of *C. grandiflora* carrying *CgS12* with pollen from *C. orientalis* results in pollen tube growth (compatible reaction), demonstrating that *C. orientalis* SCR is not functional.

### A conserved S-linked sRNA is associated with dominant expression of C. orientalis SCR

Under most circumstances, loss of function mutations are predicted to be recessive, as a single copy of a functional allele is generally sufficient to result in a complete phenotype (Kacser and Burns 1981). Here, SC is associated with a frameshift deletion at *SCR*, yet it is dominant in our F2s. Hence, we investigated whether the small RNA-based mechanism that governs dominance hierarchies among *S*-alleles in *Arabidopsis* (Durand et al. 2014) could also explain dominance of SC in our case. Specifically, if the *C. orientalis S*-haplotype encodes a trans-acting sRNA that represses expression of *C. grandiflora SCR* in *S*-locus heterozygotes, SC could be dominant even if it is due to a loss of function mutation in *C. orientalis SCR*.

In *A. halleri S12*, an *S*-linked sRNA-based dominance modifier termed *Ah12mirS3* has been identified (Durand et al. 2014), and we found the corresponding *mirS3* sRNA precursor region to be conserved (91.3% sequence identity) in *C. orientalis* (fig. 4A, fig. S1, Supplementary Material). To assess whether expression of *C. orientalis Ah12mirS3-like* sRNA (*ComirS3*) was associated with repression of the *C. grandiflora SCR* allele passed on in our cross through the F1 plant, we sequenced and assembled the *C. grandiflora S*-haplotype segregating in our F2 population, and analyzed *SCR* and sRNA expression in flower buds of 19 F2s. We detected expression of *ComirS3* sRNAs (fig. 4A) in F2s harboring the *C. orientalis S*-haplotype, but not in *C. grandiflora S*-homozygotes (fig. 4B). The most abundant *ComirS3* sRNA was highly similar to the *Ah12mirS3* sRNA and had a predicted target within the intron of *C. grandiflora SCR* allele (fig. 4D) with similar sRNA-target affinity as for functional *Arabidopsis* dominance modifiers (Durand et al. 2014, Burghgraeve et al. 2018). As expected if *ComirS3* sRNAs silence *C. grandiflora SCR, C. grandiflora SCR* was specifically downregulated in *S*-locus heterozygotes (fig. 4C). These results are consistent with *S*-linked sRNAs conferring dominance of the SC *C. orientalis S*-haplotype.

**Figure 4.**
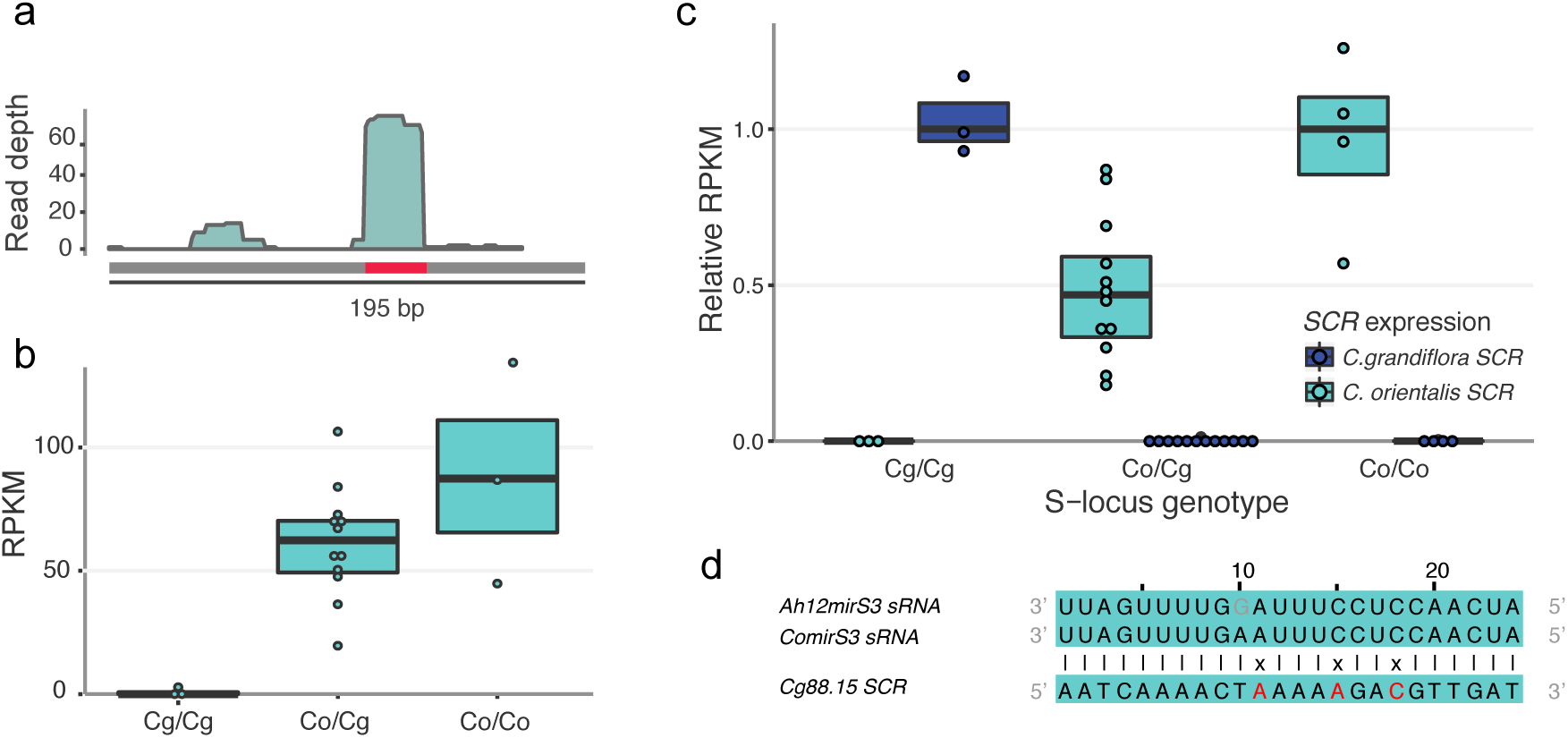
A conserved, *S*-linked *C. orientalis* sRNA is associated with repression of *C. grandiflora SCR* in *S*-locus heterozygotes. **a.***C. orientalis* expresses *S*-linked small RNAs (sRNAs) homologous to *A. halleri S12 Ah12mirS3* in flower buds. The location of *Ah12mirS3* expressed in *A. halleri S12* is indicated in red, and the grey box indicates the length of the sRNA precursor region. **b.**Expression (reads per kilobase of transcript per million mapped reads, RPKM) of 18-27 nt sRNAs in the *Ah12mirS3*-like RNA precursor region in flower buds differs between F2s with different *S*-locus genotypes (Kruskal-Wallis χ^2^=7.830, P=0.012): “Cg/Cg” and “Co/Co” are homozygous for the *C. grandiflora* or *C. orientalis S*-allele respectively, wheras “Co/Cg” are heterozygous. Only homozygotes or heterozygotes for the *C. orientalis S*-allele express sRNAs in the *Ah12mirS3*-like RNA precursor region (Dunn’s test P<0.01 for both comparisons Cg/Cg vs. Co/Cg and Cg/Cg vs. Co/Co). **c.**Relative expression (RPKM) of *C. grandiflora SCR* (blue) and *C. orientalis SCR* (turquoise) in F2 individuals with different *S*-locus genotypes, labeled as in b. *C. grandiflora SCR* is repressed in *C. grandiflora/C. orientalis* heterozygotes (Kruskal-Wallis χ^2^(2) = 9.9383, *P* < 0.01, Dunn’s test Z(2) = 2.25, *P* = 0.012 for Co/Cg vs Cg/Cg). Values for *C. grandiflora* are relative to the median RPKM of *C. grandiflora* homozygotes, whereas those for *C. orientalis SCR* are relative to the median RPKM of *C. orientalis* homozygotes. **d.***mirS3* 24-nt small RNA sequences of *A. halleri S12 (Ah12mirS3)* and *C. orientalis (ComirS3)* and the predicted target in *C. grandiflora* Cg88.15 *SCR*, located 665 bp from exon 1 and 183 bp from exon 2.

### Timing of loss of self-incompatibility in C. orientalis

After the loss of SI, the *S*-locus is expected to evolve neutrally and polymorphism at the *S*-locus can be used to estimate the timing of loss of SI (Guo et al. 2009). We analyzed 38 full-length *S*-locus sequences and estimated an upper bound for the timing of loss of SI in *C. orientalis* as the time to the most recent common ancestor (TMRCA) of *C. orientalis, C. bursa-pastoris* B and *C. grandiflora CgS12 S*-haplotypes. As *C. orientalis* was selfing when it contributed to the origin of *C. bursa-pastoris* (Douglas et al. 2015), we obtained a lower bound as the TMRCA of *C. orientalis* and *C. bursa-pastoris* B *S*-haplotypes (fig. 5). Based on these analyses, we infer a loss of SI in *C. orientalis* between 2.6 Mya and 70 kya (fig. 5, table S5, Supplementary Material) under a exponential population size change model. Very similar estimates were obtained under a constant population size model (table S5, Supplementary Material), suggesting that these results are robust to assumptions regarding population size changes.

**Figure 5.**
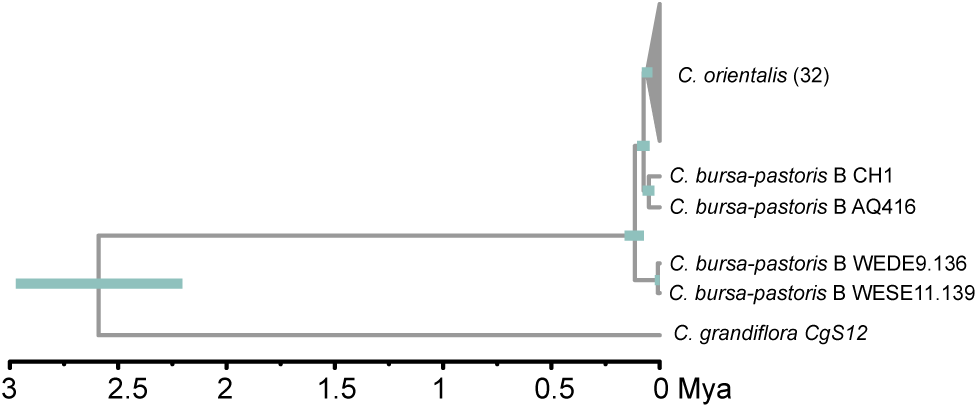
The timing of loss of self-incompatibility in *C. orientalis*. Phylogenetic tree showing relationships among *S*-haplotypes and estimates of the timing of the loss of self-incompatibility (SI) in *C. orientalis* based on analyses in BEAST2. Green bars at nodes indicate 95% credible intervals of the time to the most recent common ancestor (TMRCA). The TMRCA of *C. grandiflora CgS12* and *C. orientalis* + *C. bursa-pastoris* B represents an upper bound for the timing of loss of SI in *C. orientalis*. Because *C. orientalis* was selfing when it contributed to the origin of *C. bursa-pastoris*, the TMRCA of *C. orientalis* and *C. bursa-pastoris* B represents a lower bound on the timing of loss of SI.

## Discussion

Here, we show that loss of SI in *C. orientalis* maps as a dominant trait to the *S*-locus. We identify a frameshift deletion in the male specificity gene *SCR*, confirm loss of male SI specificity, and identify a conserved sRNA that could be responsible for dominance of SC. Our results are consistent with theory predicting a role for *S*-linked mutations in the loss of SI (Porcher and Lande 2005), and suggest that mutations in the male specificity component were important for degeneration of SI. Our finding that SC is dominant agrees with Haldane’s prediction that dominant alleles enjoy a higher fixation probability in outcrossers (Haldane 1927).

The *C. orientalis SCR* deletion that we identified by comparing these *S*-haplotypes is expected to lead to the loss of 5 of 8 conserved cysteine residues in the SCR protein, and could thus be expected to lead to the loss of male specificity. The *SCR* deletion was fixed in a broad sample of *C. orientalis*, as we would expect if it arose early during the transition to selfing, and it was also found in the allopolyploid *C. bursa-pastoris*, suggesting that the shift to SC in *C. orientalis* predated the origin of *C. bursa-pastoris*.

Theory predicts that mutations that disrupt male SI specificity should be more strongly selected for during the transition to selfing (Uyenoyama et al. 2001; Busch and Schoen 2008; Tsuchimatsu and Shimizu 2013). Indeed, mutations that disrupt male SI specificity should have an advantage both when spreading through seeds and pollen, because they avoid recognition and rejection when they spread through outcross pollen, in contrast to mutations that disrupt female specificity, which only have an advantage when there is pollen limitation (Uyenoyama et al. 2001; Busch and Schoen 2008; Tsuchimatsu and Shimizu 2013). It is possible that this advantage contributed to the spread of the SC mutation in *C. orientalis*, as has been hypothesized for the loss of SI through decay of male SI specificity in *L. alabamica* (Busch et al. 2011), European accessions of *A. thaliana* (Tsuchimatsu et al. 2010) and *A. kamchatica* (Tsuchimatsu et al. 2012).

Through crosses between *C. orientalis* and *C. grandiflora* individuals harboring highly similar *S*-haplotypes, we functionally confirmed that male SI specificity was indeed lost in *C. orientalis*, as the pollen of *C. orientalis* germinated on the stigma of individuals harboring the highly similar but functional *CgS12* haplotype. However, we cannot strictly rule out a contribution of *S*-linked mutations that disrupt female SI specificity, as controlled crosses indicated that female SI specificity was also impaired in *C. orientalis*. This finding illustrates a general challenge for studies that aim to identify causal changes for the loss of SI - after SI has been lost, the *S*-locus is expected to evolve neutrally and additional mutations that impair the function of *S*-locus genes can accumulate without cost (barring pleiotropic constraints). In this study, we did not find major-effect mutations in *SRK* or *S*-linked modifier loci in *C. orientalis*, but we cannot currently rule out that subtle changes to the sequence or expression of these genes, perhaps accumulating after the initial loss of SI, have affected their function. Such secondary decay at the *S*-locus is expected to become more likely over time after the loss of SI. To test this hypothesis, transformation experiments will now be required.

Here, we estimate that the loss of SI in *C. orientalis* occurred less than 2.6 Mya but before 70 kya, which means that loss of SI could have occurred farther back in time in *C. orientalis* than in the selfing diploid *C. rubella* as well as other well-studied cases in the Brassicaceae (e.g. ~50-100 kya in *C. rubella*, Slotte et al. 2013; ~12-48 kya in the a2 race of *L. alabamica*, Busch et al. 2011). In comparison to the recently derived selfer *C. rubella, C. orientalis* has strongly reduced genome-wide polymorphism levels (Douglas et al. 2015), shows increased reproductive isolation through endosperm development defects in crosses to *C. grandiflora* (Lafon-Placette et al. 2018), and possibly exhibits a lower genomic content of transposable elements (Ågren et al. 2014). An older origin of selfing in *C. orientalis* than in *C. rubella* would be compatible with these findings, as selfing is expected to result in reduced polymorphism genome-wide and affect TE content (Wright et al. 2013; Slotte 2014). While the shift to selfing was clearly independent in *C. orientalis* and *C. rubella*, which harbor different *S*-haplotypes (fig. 2A), both shifts involved fixation of a single *S*-haplotype (Guo et al. 2009; Slotte et al. 2012), in contrast to the situation in *A. thaliana*, where multiple *S*-haplogroups are still segregating (Durvasula et al. 2017; Tsuchimatsu et al. 2017).

Population geneticists have long predicted that dominant beneficial mutations should have a higher fixation probability than recessive ones (Haldane 1927), a phenomenon termed “Haldane’s sieve”. Our finding that SC is dominant over SI is consistent with this prediction, and agrees with results for several other wild Brassicaceae species (e.g. *L. alabamica;* Busch et al. 2011; *A. kamchatica;* Tsuchimatsu et al. 2012; *C. rubella;* Slotte et al. 2012; but see Mable et al. 2017 for an example of a recessive loss of SI in *A. lyrata)*. Our results further suggest that a small RNA-based mechanism could explain dominance of SC. If this is the case, the dominance of the SC phenotype will depend on the exact combination of *S*-alleles and their position in the dominance hierarchy. Interestingly, in both *C. orientalis* and *C. rubella*, SC is linked to relatively dominant *S*-haplotypes. Taken together, these findings suggest that dominant SC mutations have an advantage over recessive mutations, at least early during the transition to selfing, and that the lower population frequencies or higher *S*-linked load (Llaurens et al. 2009) of dominant *S*-alleles do not prevent mutations in such alleles from contributing to recurrent loss of SI.

## Materials and Methods

### Plant material and growth conditions

We surface-sterilized seeds of *C. orientalis, C. bursa-pastoris* and *C. grandiflora* accessions (supplementary table S1, Supplementary Material), plated them on ½ MS medium (Murashige and Skoog basal salt mixture, Sigma-Aldrich Co. MI, USA) and stratified the seeds at 2-4°C in the dark for two weeks. Plates were then moved to climate chambers (16 h light at 20°C / 8 h dark at 18 °C, 70 % maximum humidity, 122 uE light intensity) to germinate. After one week, seedlings were transplanted to soil in pots. For genotyping and whole-genome resequencing, leaf samples for DNA extractions were collected from >3 week old plants and dried in silica gel. For bacterial artificial chromosome (BAC) library construction, leaf samples were collected after 48 h dark treatment and were immediately flash-frozen in liquid N_2_. For RNA extractions, mixed-stage floral buds and leaf samples were collected in the middle of the light period and immediately flash-frozen in liquid N_2_.

### Genetic mapping of loss of SI in C. orientalis

We generated an interspecific *C. orientalis* × *C. grandiflora* F2 mapping population which segregated for SI/SC by crossing *C. orientalis* accession Co2008-1 as seed parent to *C. grandiflora* accession Cg88.15 as pollen donor (supplementary table S1, Supplementary Material). Because *C. orientalis* × *C. grandiflora* F1 seeds were aborted prior to full development, generating viable F1 seeds required embryo rescue (for details, see supplementary text, Supplementary Material). F1 individuals were SC, and we collected F2 seeds from one autonomously self-pollinated F1 individual. Our final mapping population consisted of a total of 350 F2 individuals. We extracted DNA from all F2 individuals using a Qiagen DNeasy kit (Qiagen, Venlo, The Netherlands) and genotyped them at 998 SNPs at SciLifelab Stockholm (for details, see Supplementary Material).

We scored SI/SC in a total of 321 F2 individuals. SI/SC was visually scored as presence or absence of silique formation on mature individuals. In addition, we assessed the success of 3-6 manual self-pollinations for 204 F2 individuals. In the case of a discrepancy between seed set after manual self-pollination and silique formation after autonomous self-pollination, we used the scoring based on manual self-pollination. To validate that the SI phenotype was due to pollen tube growth arrest and the lack of seed development following self-pollination was not due to e.g. inbreeding depression or later-acting genetic incompatibilities, we assessed pollen tube growth in the pistil after manual self-pollination in a subset of 10 F2 individuals scored as SI (supplementary text, Supplementary Material).

We generated a linkage map and mapped quantitative trait loci (QTL) for SI/SC status in R/Qtl (Broman et al. 2003). The final linkage map had 549 SNPs after removal of SNPs with redundant genotype information or that showed segregation distortion. We mapped QTL for SI/SC, encoded as a binary trait, using interval mapping and the Haley & Knott regression method (Haley and Knott 1992) in intervals of 1 cM. A 1% genome-wide significance threshold was obtained by 1000 permutations and we estimated credible intervals of significant QTL as 1.5-LOD drop intervals. We estimated the additive allelic effect and dominance deviation at significant QTL using the R/Qtl effectscan function.

### Sequencing, assembly and annotation of the S-locus in Capsella

To identify putative causal genetic changes responsible for loss of SI in *C. orientalis*, we conducted targeted sequencing and assembly of *S*-haplotypes by long-read sequencing of bacterial artificial chromosome (BAC) clones containing the *S*-locus, as previously described (Bachmann et al. 2018) (see also supplementary text, Supplementary Material). We generated BAC libraries and conducted targeted long-read sequencing and assembly of *S*-haplotypes of two SC *C. orientalis* accessions, four SC *C. bursa-pastoris* accessions and two SI *C. grandiflora* accessions. The two *C. grandiflora S*-haplotypes presented here were chosen from a larger set of 15 *S*-haplotypes (to be fully presented elsewhere) to represent the *C. grandiflora S*-haplotype segregating in our F2 population as well as a *C. grandiflora S*-haplotype from the same haplogroup as the *S*-haplotype of *C. orientalis* (see “Phylogenetic analyses of *S*-locus sequences” below; supplementary table S1, Supplementary Material). In total, we here present eight full-length *S*-locus haplotypes obtained by targeted long-read sequencing (supplementary tables S1 and S2, Supplementary Material).

BAC clones were sequenced to high coverage (150-400x) using PacBio SMRT sequencing (supplementary table S2, Supplementary Material). Short-read sequencing data for all BACs were generated on an Illumina MiSeq (>380 x; supplementary table S2, Supplementary Material) and used for indel error correction as described previously (Bachmann et al. 2018). All sequencing was done at the SciLifeLab NGI in Uppsala, Sweden. Sequences were assembled in HGAP3.0 (Chin et al. 2013), except for the *S*-haplotype of Cg88.15, for which Canu v.1.7 (Koren et al. 2017) was used instead as HGAP3.0 assembly was unsuccessful.

We annotated our *S*-locus assemblies as previously described (Bachmann et al. 2018). Briefly, we used Augustus v3.2.3 (Stanke et al. 2004) and RepeatMasker v4.0.7; http://www.repeatmasker.org), run via Maker v2.31.9 (Holt and Yandell 2011) with *Arabidopsis thaliana* as a model prediction species and using protein homology data for *SRK, U-box* and *ARK3* from *Arabidopsis lyrata* and *Arabidopsis halleri*. Due to the high levels of sequence diversity at the key *S*-locus genes *SRK* and *SCR*, they were difficult to annotate automatically. Sequence similarity (BLASTN) to known *SRK* exon 1 sequences was used to accept candidate loci as *SRK*, while we used close similarity to *ARK3* as a rejection criterion. To annotate *SCR*, we used a window-based approach to screen for the characteristic pattern of cysteine residues after translation of the DNA sequence in all three frames, as described previously (Bachmann et al. 2018). Using this approach, we identified a region highly similar to *A. halleri SCR* in *S*-locus haplotype *S12* (GenBank accession number KJ772374.1) in our *C. orientalis S*-locus BAC sequences. Using BLASTN we found high similarity between the Cg88.15 *S*-haplotype segregating in our F2 population and the *S*-haplotype *A. halleri AhS4* (GenBank accession KJ461484), and the Cg88.15 *S*-haplotype was therefore annotated by reference to the *A. halleri AhS4* sequence annotation.

### Phylogenetic analyses of S-locus sequences

Using a dataset of Brassicaceae *SRK* exon 1 and *ARK3* sequences downloaded from Genbank for a previous study (Bachmann et al. 2018) we generated an alignment of *SRK* exon 1 sequences using the MAFFT v7.245 & E-INS-I algorithm (Katoh et al. 2002) with manual curation and error correction in SeaView v4.6 (Gouy et al. 2010). We generated a maximum likelihood phylogenetic tree from the alignment of *SRK* sequences with RaXMl v8.2.3 (GTRGAMMA model and 1000 bootstraps replicates) and then plotted the *SRK* phylogeny in R v. 3.3.1 (R Core Team 2017). In this phylogeny, the *C. grandiflora* Cg2-2 *S*-haplotype clustered with the *S*-haplotypes of *C. orientalis* and the *C. orientalis-derived* subgenome of *C. bursa-pastoris* (i.e. the *C. bursa-pastoris* B subgenome). Due to the high sequence similarity (93.4% protein sequence identity at *SRK*) of the Cg2-2 *C. grandiflora S*-haplotype to *A. halleri S12* (GenBank accession number KJ772374.1) we termed this *S*-haplotype *CgS12*. We further assessed sequence conservation across the entire ~100 kbp *S*-locus by aligning *S*-locus sequences using LASTZ v1.03.54 (Harris 2007) and calculating pairwise sequence conservation in 250 bp sliding windows.

### Candidate mutations for the loss of SI in C. orientalis

To identify candidate causal mutations for the loss of SI in *C. orientalis*, we analyzed sequence alignments of the two key *S*-locus genes *SRK* and *SCR*, as well as of the *S*-linked *U-box* gene, which may act as a modifier of the SI response (Liu et al. 2007). Specifically, we searched for major-effect variants such as frameshifts, premature stop codons or non-consensus splice sites present in sequences from the SC *C. orientalis* and/or in the SC *C. bursa-pastoris* B subgenome, which is derived from *C. orientalis* (Douglas et al. 2015), but not in sequences from the same haplogroup found in the SI species *C. grandiflora* and *A. halleri* (i.e. the *C. grandiflora CgS12* and *A. halleri S12* haplotypes).

### Bioinformatic processing of RNAseq data

RNAseq data was trimmed with Trimmomatic v.0.36 (Bolger et al. 2014) and reads were mapped using STAR v.2.2.1 (Dobin et al. 2013). For small RNA sequencing reads, we mapped reads of length 18-27 nt using STAR v.2.2.1 (Dobin et al. 2013). Expression was quantified as RPKM (the number of reads per kb per million mapped reads; Mortazavi et al. 2008).

### Expression of S-locus genes in C. orientalis

To assess whether *SRK, SCR* and *U-box* were expressed in *C. orientalis* flower buds, we generated RNAseq data from mixed-stage flower buds of two *C. orientalis* accessions (Co1719/11 and Co1979/09; table S1, Supplementary Material) as previously described (Steige et al. 2017). For comparison, we also generated RNAseq data from leaf samples from the same individuals. Reads were processed as described in “Bioinformatic processing of RNAseq data” above, and mapped to a modified v1.0 reference *C. rubella* assembly (Slotte et al. 2013), where the *S*-locus region (scaffold_7 7523601:7562919) was masked and our *S*-locus assembly from *C. orientalis* Co1719/11 was added. We also conducted qualitative RT-PCR with specific primers to *SCR* in *C. orientalis* and *C. grandiflora CgS12*, to assess the expression of *SCR* in flower buds of both the Co1719/11 and Co1979/09 accessions, as well as in three *C. grandiflora* individuals harboring *CgS12* (supplementary fig. S5, Supplementary Material).

### Assessing the functionality of C. orientalis SCR by interspecific crosses

We performed controlled crosses to verify that *C. grandiflora CgS12* conferred SI, and to assess the functionality of SCR in *C. orientalis*. To verify functional SI in *C. grandiflora* carrying *CgS12*, we performed 12 manual self-pollinations of a *C. grandiflora* individual carrying the *CgS12 S*-haplotype. We note that the identity of the other *S*-haplotype in this individual is unknown and we were unable to identify it using PCR-based screening. However, we were able to verify expression of *CgSCR12*, indicating that the other *S*-allele is not dominant over *CgS12* at the pollen level. We further assessed the success of manual self-pollination of *C. orientalis* by performing 6 manual self-pollinations. To assess whether *C. orientalis* SCR is functional, we crossed *C. grandiflora* harboring *CgS12* as a seed parent to *C. orientalis* as a pollen donor. We performed a total of 112 crosses of this type, with two different *C. orientalis* accessions as pollen donors and three different CgS12-carrying *C. grandiflora* individuals as seed parents (supplementary table S1, Supplementary Material). If *C. orientalis* SCR is functional, and provided that *CgS12 SRK* is expressed, then we expect this cross to be incompatible, whereas if *C. orientalis* SCR is nonfunctional, the cross should be compatible. The reciprocal cross of the same individuals was also carried out with the same accessions (total 84 crosses of this type), to test whether female SI specificity is functional in *C. orientalis*. Finally, we performed 12 crosses of *C. grandiflora* harboring other *S*-haplotypes to *C. grandiflora* harboring *CgS12*, and 12 to *C. orientalis*. These crosses are expected to be successful. We observed pollen tube growth in the pistil 12 hours after pollination. Pistils were fixed in EtOH: acetic acid 9:1 for > 2 hours, softened in 1N NaOH 60°C for 20 minutes and stained with 0.01% decolorised aniline blue in 2% solution of K_3_P0_4_ for 2 hours. Pollen tubes were visualised by mounting the pistils on a microscope slide which was examined under an epifluorescence microscope (Zeiss Axiovert 200M). We compared the number of pollen tubes among different types of crosses using a Kruskal-Wallis test (supplementary fig. S4, Supplementary Material).

### The role of small RNA-based dominance modifiers for dominance of SC in C. orientalis

To test whether the dominant expression of SC in our F2s (see Results) could be mediated by small RNA-based dominance modifiers, we conducted additional sequence and expression analyses. First, we identified a region in our *C. orientalis S*-haplotypes with high sequence similarity (91.3%) to the *A. halleri S12* small RNA precursor *Ah12mirS3* from (Durand et al. 2014). We generated small RNA and RNA sequencing data from flower buds of 19 F2s, representing all three *S*-locus genotypes in our F2 mapping population (12 heterozygotes, 4 and 3 individuals homozygous for the *C. orientalis* or the *C. grandiflora S*-haplotype, respectively). Reads were processed as described in “Bioinformatic processing of RNAseq data” above, and mapped to a modified v1.0 reference *C. rubella* assembly (Slotte et al. 2013), where the *S*-locus region (scaffold_7 7523601:7562919) was masked and the *S*-haplotype of *C. orientalis* Co1719/11 was added. We quantified expression of sRNAs in the *Ah12mirS3*-like sRNA precursor region, hereafter termed *ComirS3* sRNAs, and compared expression in the three genotypes to test whether small RNAs in this genomic region were expressed specifically in F2s with a *C. orientalis S*-allele.

To test whether *C. grandiflora SCR* was repressed in heterozygous F2s we quantified the expression of *C. orientalis* and *C. grandiflora SCR* in our F2s. We mapped F2 RNAseq reads from flower buds to a modified *C. rubella* reference containing both the Co1719/11 *S*-haplotype and the *C. grandiflora* Cg88.15 *S*-haplotype segregating in our F2 population, and quantified the expression of *C. orientalis* and *C. grandiflora SCR* in all three genotypes, respectively.

To identify targets of *ComirS3* small RNAs we took all expressed small RNA (18-27 nt) in flower buds samples from three F2 individuals homozygous for the *C. orientalis S*-haplotype and searched for small RNA targets within 1 kb of *SCR* of the *C. grandiflora* Cg88.15 *S*-haplotype. Small RNA targets were identified using a Smith & Waterman algorithm (Smith and Waterman 1981) with scoring matrix: match=01, mismatch=−1, gap=−2, G:U wobble=−0.5 as previously described (Durand et al. 2014).

### Population genomic analyses

To assess whether the *SCR* deletion at the *S*-locus was fixed in *C. orientalis*, we analyzed whole-genome resequencing data from additional *C. orientalis* accessions, in total covering 30 accessions from 18 populations (table S1, Supplementary Material). We mapped trimmed data to a *C. rubella* reference modified to include the *C. orientalis* haplotype of accession Co1719/11 using BWA-MEM (Li 2013) and called variants using GATK 3.8 (McKenna et al. 2010; DePristo et al. 2011; Van der Auwera et al. 2013) HaplotypeCaller using the GVCF mode to call all sites. We filtered sites following GATK recommended hard filtering with the following parameters; QD < 2.0 || FS > 60.0 || MQ < 40.0 || MQRankSum < −12.5 || ReadPosRankSum < −8.0. We required a minimum read depth of 15 and a maximum of 200. Finally, we scored the presence or absence of the *SCR* deletion in our samples. Because *C. orientalis* is highly homozygous, self-compatible, and has low levels of polymorphism genome-wide (Douglas et al. 2015), this approach is expected to work well, as long as a *C. orientalis S*-haplotype is included in the reference genome.

We used a strategy similar to that in (Guo et al. 2009) to estimate a lower and upper bound of the timing of the loss of SI in *C. orientalis*. We obtained a lower bound for the timing of the loss of SI by estimating the time to the most recent common ancestor (TMRCA) based on full-length *C. orientalis* and *C. bursa-pastoris* B *S*-locus sequences. This is possible because genome-wide haplotype sharing between *C. orientalis* and the *C. bursa-pastoris* B subgenome (Douglas et al. 2015), indicates that the ancestor of *C. orientalis* that contributed to formation of *C. bursa-pastoris* was already selfing. Therefore, including *C. bursa-pastoris* B sequences can allow us to increase the precision of our estimates of the lower bound. To obtain an upper bound for the timing of the loss of SI we estimated the TMRCA for *C. orientalis, C. bursa-pastoris* B and *C. grandiflora CgS12*.

For analyses of the timing of loss of SI, our final alignment contained 37 *S*-locus sequences including the *C. grandiflora* ancestral *S*-haplotype (*CgS12*), 4 *C. bursa-pastoris* subgenome B *S*-haplotypes and *S*-haplotype data for 32 *C. orientalis* individuals (supplementary text, Supplementary Material). Sequences were aligned using block alignment using Muscle v.3.8.31 (Edgar 2004) as implemented in AliView v.1.20 (Larsson 2014). The total length of the *S*-locus alignment was 33,485 bp, 22,689 bp had indels in at least one sequence, 9,835 sites were invariant and 876 sites were polymorphic. The alignment was partitioned into coding and non-coding regions and sites with indels and missing data were pruned in further analysis.

We estimated the timing of the splits between *C. grandiflora, C. bursa-pastoris* and *C. orientalis* as well as the crown age of *C. orientalis* using a strict molecular clock in a Bayesian framework as implemented in BEAST2 (Bouckaert et al. 2014). We used a fixed clock rate assuming a mutation rate of 7×10^−9^ substitutions per sites per generation (Ossowski et al. 2010) and a generation time of one year. The best substitution models inferred in PartitionFinder2 v.2.1.1 (Lanfear et al. 2012; Lanfear et al. 2017) for the coding and non-coding partition were GTR + G and HKY + I respectively. We ran both a complex model with exponential changes in population size and a model with a constant population size, and assessed whether the more complex model gave a significant improvement in likelihood using aicm (Baele et al 2012) (table S5, Supplementary Material). We ran two chains of 10 millions generations sampled every 1000 generations and checked the convergence by visual inspection of the log-likelihood profile and assuring ESS value above 200. The posterior distribution of trees was used to build a maximum clade credibility tree and estimate node age and 95% confidence interval using TreeAnnotator (Drummond et al. 2012).

## Acknowledgements

We thank Timothy Paape for helpful discussion, Daniel Koenig and Detlef Weigel for having made *C. orientalis* resequencing data publicly available, Cindy Canton for help with plant care and sampling, and Christian Tellgren-Roth for help with BAC assembly. The authors acknowledge support from the National Genomics Infrastructure (NGI) / Uppsala Genome Center / SNP&SEQ Technology Platform and UPPMAX for providing assistance in massive parallel sequencing and computational infrastructure. Work performed at Uppsala Genome Center has been funded by RFI / VR and Science for Life Laboratory, Sweden. The SNP&SEQ Platform is also supported by the Swedish Research Council and the Knut and Alice Wallenberg Foundation. V.C. acknowledges support by a grant from the European Research Council (NOVEL project, grant #648321). The authors thank the French Ministère de l’Enseignement Supérieur et de la Recherche, the Hauts de France Region and the European Funds for Regional Economical Development for their financial support to this project. This work was supported by a grant from the Swedish Research Council (grant #D0432001) to T.S.

## Author contributions

T.S. designed the experiments. J.B., A.T., C.L.-P., K.A.S., C.C. and W.M. performed the experiments, J.B., A.T. and A.D. generated the data. J.B. analyzed sRNA expression and targets, A.T. analyzed and annotated *S*-locus BACs, B.L. analyzed and annotated *S*-locus BACs and performed BEAST analyses, M.F. analyzed QTL mapping and expression data, B.N. contributed reagents/materials/analysis tools, and A.D. generated full-length *S*-locus alignments. All authors contributed to the writing of the paper.

